# *synapsis*: A Bioconductor package to automate the analysis of meiotic double-strand break repair and crossover formation

**DOI:** 10.1101/2022.07.14.500141

**Authors:** Lucy McNeill, Vanessa Tsui, Wayne Crismani

## Abstract

**Summary:** Immunofluorescent staining is commonly used to generate images to characterise cytological phenotypes. The manual quantification of DNA double-strand breaks and their repair intermediates during meiosis using image data requires a series of subjective steps, from image selection to the counting of particular events per nucleus. Here we describe *synapsis*, a Bioconductor package, which includes a set of functions to automate the process of identifying meiotic nuclei and quantifying key double-strand break formation and repair events in a rapid, scalable and reproducible workflow, and compare it to manual user quantification. The software can be extended for other applications in meiosis research, such as incorporating machine learning approaches to categorise meiotic substages.

**Availability and implementation:** *synapsis* can be freely downloaded and installed at: http://bioconductor.org/packages/release/bioc/html/synapsis.html

R functions and further information can be found at https://gitlab.svi.edu.au/drr-public/synapsis

## 1 Introduction

Meiosis is a special type of cell division that is necessary to make sperm and eggs ^6^. During meiosis chromosomes are recombined and their numbers halved through highly regulated processes. Two key steps in the meiotic process are studied heavily and can be visualised with immunofluorescent (IF) techniques: 1) the reorganisation of chromosome structure is unique at meiosis because homologous chromosomes pair and align in a process known as synapsis, and this can be appreciated when visualising the synaptonemal complex (Fig. 1a) ^2,4,11^; 2) DNA double-strand breaks (DSBs) form in high numbers and are repaired by meiosis-specific recombination pathways. This is a homeostatic process, which lends itself to visualisation as many of the proteins involved form foci ^3,5,7,10^. Most meiotic immunofluorescent localisation analyses currently rely on subjective manual counting or ImageJ macros, the latter requiring significant user intervention to run. To the best of our knowledge this is the first open access software package designed for the analysis of meiotic immunofluorescent image data that has been optimised with samples from different taxonomic kingdoms.

**Figure 1:**
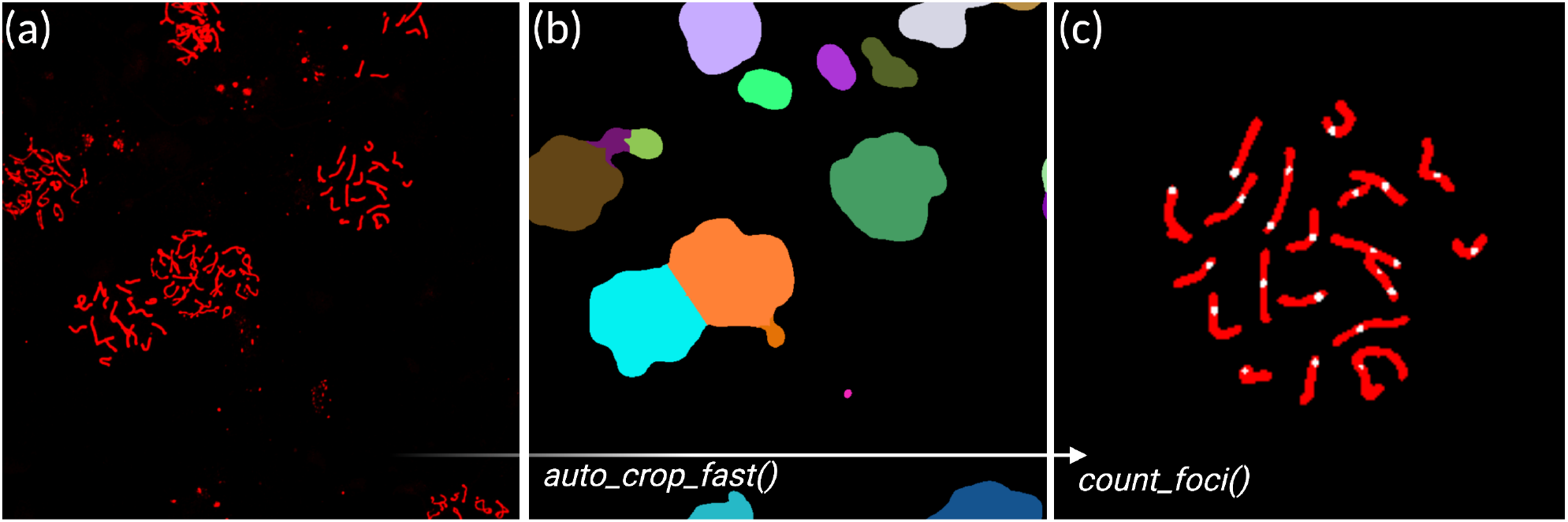
Workflow of the synapsis package. **(a)** The function *auto* crop *fast* is called on the path that corresponds to the directory which contains the images. False-coloured anti-SYCP3 signal is shown. **(b)** Watershed output of an intermediate step in the auto crop fast function. **(c)** False-colour merge of anti-MLH3 foci (white) and anti-SYCP3 masks (red) identified by the *count foci* function. Biorender was used to annotate the figure.

## 2 Usage and performance compared to other methods

*synapsis* is an open source R package whose functions are structured and documented in line with Bioconductor standards, and makes significant use of EBImage functions ^8^. The general workflow and use of functions is shown in Figure 1. *synapsis* has been optimised to analyse grayscale images from up to three channels, which have been generated with commonly used meiotic chromosome spreading techniques ^1,9^. First, individual nuclei are identified, based on markers of the synaptonemal complex, and a new folder of cropped nuclei is created. Then, in an optional step, pachytene nuclei can be identified and pixel information about their synaptonemal complexes are stored in a dataframe. Next, foci that co-localise with the synaptonemal complex are quantified along with other quantitative pixel information (e.g. foci size and intensity), which can be stored in a dataframe. Finally, a quantitative rating of the confidence in the foci counted in a given nucleus is returned. In a sense, the *synapsis* workflow runs in the opposite direction to manual user counting, where in *synapsis*, all foci that colocalise with the synaptonemal complex are quantified regardless of the meiotic substage or the signal-to-background ratio of the foci. Finally, a nucleus is assigned a suggested *keep* or *discard* label. These automated decisions are reproducible and rapid and will allow researchers to be more ambitious with the scale of their analyses and the image features that are considered.

### Cropping images of individual nuclei with *auto crop fast*

The cropping function uses pixel data from the synaptonemal complex channel (Fig. 1a) to identify nuclei. The function segments bright and relatively circular features that are identified through thresholding and watershed methods (Fig. 1b). The function also generates a folder, at a user-specified location, to which it saves cropped images of a given nucleus for all channels. For users with raw data from Nikon microscopes, we provide instructions on how to use the Python package *nd2reader* to convert .nd2 files into grayscale files for each channel to common file types (TIF, JPEG, PNG), with the option to specify the desired resolution.

### Selecting nuclei in pachtene with *get pachytene*

Using the synaptonemal complex channel of each nucleus candidate, objects that pass a tunable signalto-background threshold are identified. Bright and uniformly intense synaptonemal complexes, are characteristic of the pachytene phase, and only crops with a biologically relevant number of strand objects are classified as pachytene.

### Counting the number of colocalising foci with *count foci*

Using the individual cropped nuclei images, and optionally only the pachytene crops, foci masks of sufficiently high signal compared to local background are created via Gaussian smoothing. Foci that co-localise with a synaptonemal complex (Fig. 1c) have their number, pixel areas and intensity stored to a dataframe. Using the distribution of pixel areas of foci, we determine the “crispness criteria”. This gives a measure of how uniformly sized the foci are via the standard deviation of foci areas, divided by the total number identified times the mean. Unusually low foci counts compared to chromosome number will score poorly, as well as foci masks that are unevenly sized, which can be indicative of noise. We find that high values for this coefficient are a useful empirical measure that is consistent with nuclei that are included by manual counting.

### Comparison with manual counting

We compared the performance of synapsis to manual counting of MLH3 foci (Fig. 2). We show that manual counting of MLH3 foci between individuals from our team is positively correlated, but is not reproduced perfectly from one individual to the next due to the series of subjective decisions that must be made about what constitutes a focus. Focus counts between synapsis and manual user counts was positively correlated also, however of the two approaches, only synapsis is perfectly reproducible and scalable. Thus we feel it is reasonable to propose that *synapsis* can be considered as an open-source tool for improving throughput and reproducibility in quantitative analyses of meiotic IF data.

**Figure 2:**
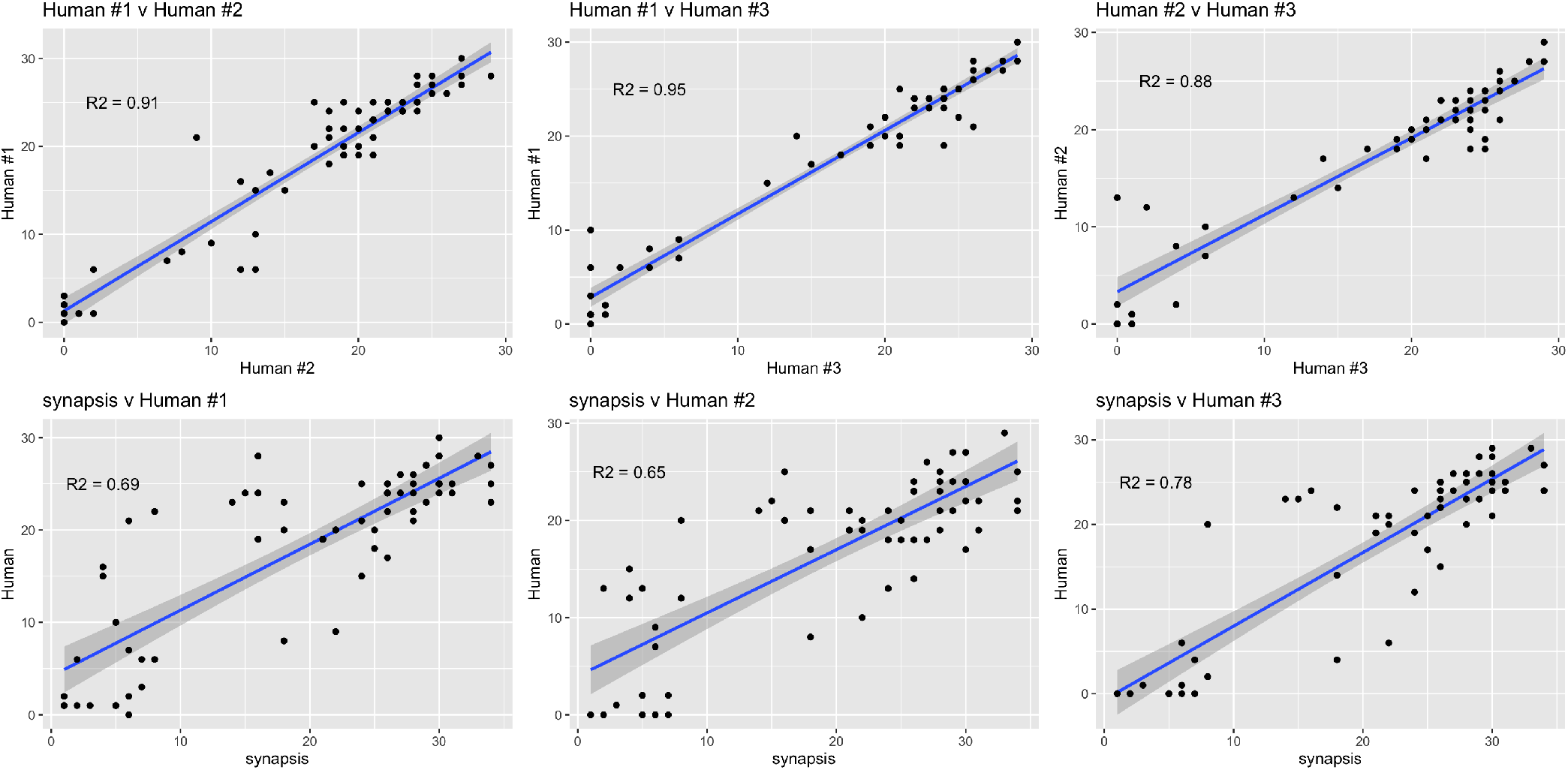
Comparison of *synapsis* against manual counting. **(a)** The function *auto crop fast* is called on the path that corresponds to the directory which contains the images. False-coloured anti-SYCP3 signal is shown. **(b)** Watershed output of an intermediate step in the auto crop fast function. **(c)** False-colour merge of anti-MLH3 foci (white) and anti-SYCP3 masks (red) identified by the *count foci* function.

## 3 Conclusion

*synapsis* provides a reproducible workflow which can automate the quantification of immunofluorescent image analysis of meiotic DSBs, repair intermediates and crossover formation. The package automates time consuming and subjective processes and can therefore assist in making robust comparisons between groups (e.g. genotypes). The software could be extended to analyse other aspects of meiosis such as interference between foci. Further, *synapsis* creates output that is compatible with machine learning approaches for applications such as meiotic substage classification, which could facilitate the efficient use of the high volumes of data that can be generated by slide-scanning microscopes.

## 4 Materials and methods

## Ethics statement

All animal procedures were approved by the Animal Ethics Committees at St Vincent’s Hospital Melbourne and conducted in accordance with Australian NHMRC Guidelines on Ethics in Animal Experimentation.

### Meiotic chromosome surface spreads and image analysis

Meiotic chromosome spreads were prepared from F1(FVB/N x C57BL/6J) adult mouse testes (8-12 weeks) using methods that were previously described ^9^. Antibodies and dilutions used were: anti-SYCP3 (sc74569 at 1:100, Santa Cruz); anti-MLH3 (1:500, gifted by Corinne Grey, Bernard de Massy and Val’erie Borde). After primary antibodies were incubated on the slides DAPI was applied at (10 µg/mL in 1xPBS) and mounted in Dako Fluorescence mounting medium. Secondary fluorescent antibodies were: donkey anti-rabbit Alexa Fluor 488 and donkey anti-rabbit Alexa Fluor 568. Images were captured using a Nikon A1R Confocal Microscope. The same sets of images of nuclei at meiotic prophase I were used for manual quantification for MLH3 foci localised to SYCP3-stained synaptonemal complexes, by three individuals and using *synapsis* v.0.99 and the *count foci* function with options: `offset factor = 8, brush size = 5, offset px = 0.3, brush sigma = 5, annotation = “on”, stage = “pachytene”, file ext = “tif”, watershed stop= “on”, crowded foci = FALSE`.

## Code availability

Source code of the latest version of *synapsis* and other functions are publicly available at GitLab repository (https://gitlab.svi.edu.au/drr-public/synapsis) under an MIT license.

## Acknowledgements

We wish to thank Ruqian Lyu, Davis McCarthy and Jeffrey Pullin for helpful feedback during the development of the package. Biorender was used to annotate Figures.

